# INAEME: Integral Neoantigen Analysis with Entirety of Mutational Events

**DOI:** 10.1101/2023.09.28.559901

**Authors:** Vladimir Kovacevic, Ognjen Milicevic, Nevena Ilic Raicevic, Milica Kojicic, Ana Mijalkovic Lazic, Nikola Skundric, Jack DiGiovanna

## Abstract

Neoantigens are peptides presented on the surface of cancer cells that can be recognized by the immune system. Multiple novel therapeutic approaches involve the administration of neoantigens to trigger immunity-induced tumor regression. Identifying neoantigens includes a personalized approach consisting of detailed analyses of the sequenced tumor tissue and its comparison with wild type to identify somatic mutations. Altered peptides are translated from nucleotides around somatic mutations, and their binding affinity and immunogenicity need further evaluation. Still, the entire bioinformatics analysis is very complex, and accurate prediction of the neoantigen candidates represents a true challenge. Here, we present the novel, integral bioinformatic analysis workflow for neoantigen discovery, denoted INAEME (Integral Neoantigen Analysis with Entirety of Mutational Events). The workflow performs integral processing of an individual’s DNA tumor-normal and RNA tumor raw reads to output prioritized neoantigen candidates. Through conducted analysis and benchmarks, our main goal was to demonstrate the necessity of taking into account a wide scope of mutational events so far not considered in the existing solutions, including phasing of variants, influence of both somatic and germline variants, positions of all transcripts, neighboring variants, and frameshifts. The influence of each mutational event on the accuracy of predicted neoepitope candidates is tested across 300 TCGA samples from multiple cancer types, including melanoma, hepatocellular carcinoma, and lung squamous cancer. The observed loss of neoantigen nests, going from 8.45% up to 23.65%, underscores the importance of accounting for the entirety of mutational events to accurately identify robust neoantigen candidates for cancer immunotherapy and vaccine development. The adaptation of the described methods in the bioinformatics analysis minimizes the existence of false positives, which are only later discovered in a laboratory environment using expensive methods such as mass spectrometry or microscopy.

## Introduction

Every cell in our body has mechanisms to cleave discarded endogenously synthesized proteins and present some of their fragments on its surface as antigens. When these antigens arise from nonsynonymous somatic mutations, they are termed neoantigens—unique molecular signatures of cancer cells. T-cells of the immune system can recognize and eliminate neoantigen-expressing tumor cells, preventing malignant progression [1]. However, cancer cells can evade immune surveillance through diverse molecular mechanisms, enabling tumor growth and metastasis [2–4].

The concept of boosting and training the immune system resulted in multiple novel immunotherapeutic approaches that involve identifying neoantigens and using them to trigger immunity-induced tumor regression [5–8][21][22]. The identification of neoantigens relies on genomic sequencing of both tumor and normal tissues, allowing for the detection of somatic mutations that alter the amino acid sequence. Peptides spanning these mutations serve as neoepitope candidates for immune recognition. To facilitate this process, several bioinformatics workflows have been developed for neoantigen discovery using next-generation sequencing (NGS) data [9–15]. Some workflows utilise machine learning algorithms, including ImmuneMirror [35], which incorporates a balanced random forest model for neoantigen prediction and prioritization. Novel solutions such as NeoDisc [36] uncover the heterogeneous antigenic landscape linked to defects in the antigen processing and presentation machinery. The differences between solutions mostly lie in the bioinformatic tools being used and the mutational events that are considered. Mutational events include any modification of the reference genome during the application of the called variants necessary to reconstruct and obtain the exact nucleotide or protein sequence from the individual (Figure 1).

**Figure 1:**
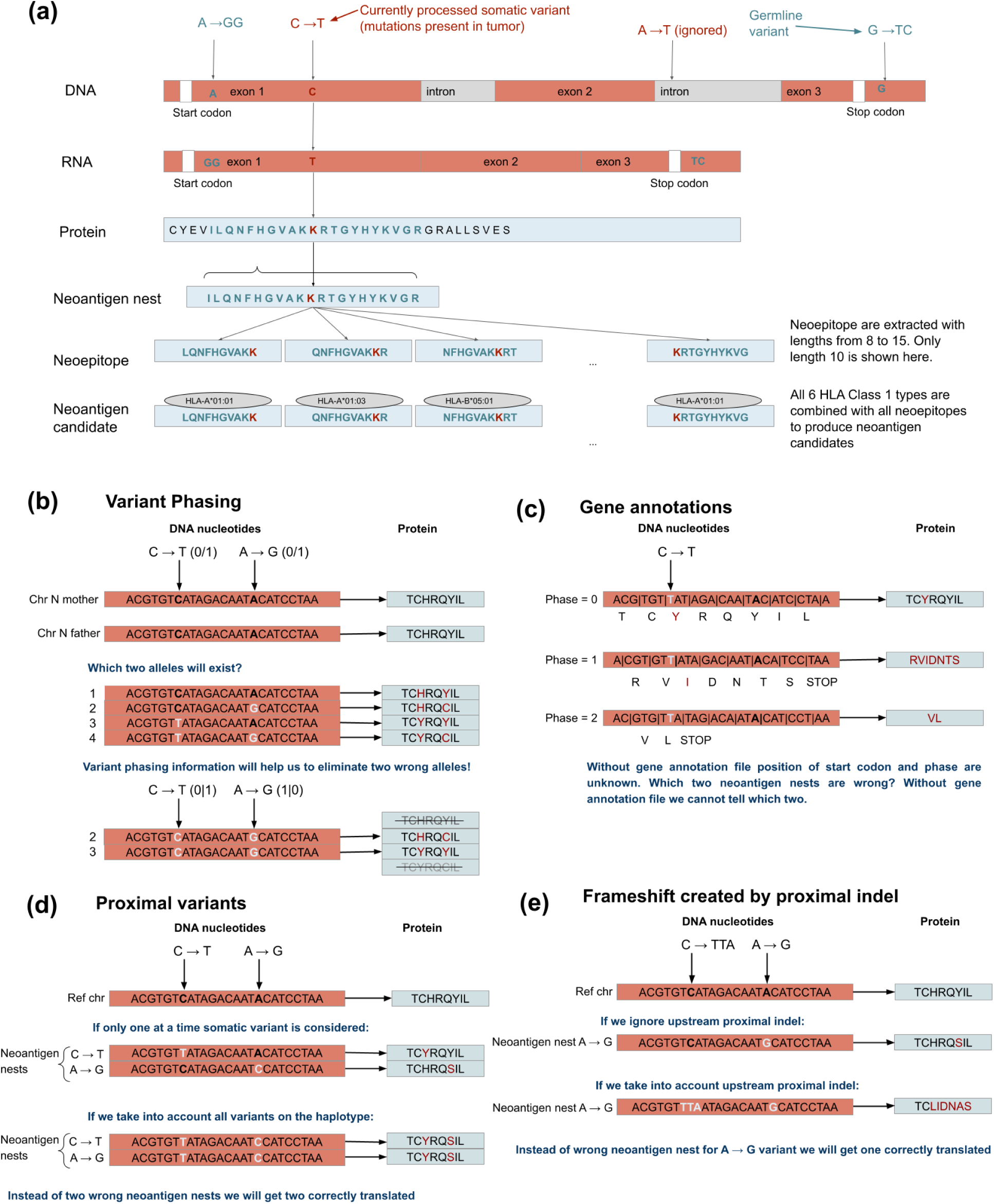
(a) Illustration of the mutational event with somatic and germline variants and their influence on protein sequence, followed by extraction of neoantigen nest from the protein, slicing neoepitopes, and creating neoantigen candidates by combining all calculated HLA types to all neoepitopes. (b) Variant phasing information enables us to determine whether variants fall on the same haplotype. If it is available it is possible to exclude nonexistent neoantigen nests. (c) Custom gene annotation file enables correct starting position and phase for translating the nucleotide sequence to protein (d). Proximal variants influence alters neoantigen nest (e). Frameshift created by proximal indel can almost completely alter the neoantigen nest.

Figure 1a shows an example of applying a variant to the reference genome to produce a neoantigen candidate. We considered only non-synonymous somatic variants in the gene’s coding regions and ignored synonymous and intron-falling variants. The DNA chain of nucleotides is transcribed to mRNA, which is, after splicing, translated to protein. For better referencing, the short protein sequence around the altered amino acid(s) will be called the neoantigen nest. In our case, the size of the neoantigen nest is 21 amino acids (10 amino acids on each side of the altered one). A peptide extracted from a neoantigen nest that contains altered amino acid(s) is called a neoepitope. A neoepitope paired with an independently calculated individual’s HLA Class 1 type forms a neoantigen candidate.

One of the first publicly available solutions was pVACseq [9]. Initially, it considered only the modifications of the reference genome from single-nucleotide variations, while later versions also considered indels, but not the influence of proximal (neighboring) variants (Figure 1d-e). pVACSeq workflow starts from already reconstructed variants and its structure to our knowledge is not optimized and described in any of the existing workflow description languages which would make it portable and reproducible [18]. pVAC does consider RNA expression, but not the RNA variant calling for determining whether the expressed transcript of the given allele is a reference or alternative (with variant). pVACSeq supports a wide variety of MHC Class I and MHC Class II binding prediction tools. Another solution, CloudNeo workflow [10], is described in Common Workflow Language (CWL). It also starts from a reconstructed variant, but does not consider RNA expression influence. Personalized genomic vaccine pipeline (PGV) [11] is an integrated solution taking into account the influence of indels, proximal variants, and RNA expression, but PGV does not support custom gene annotations and the influence of frameshifts from neighboring indels. Proximal variant correction (PVC) workflow [12] provides integral analysis with an emphasis on considering the influence of proximal variants. The authors do not take into account the influence of transcript positions and frameshifts, stating that it is substantially complicated to consider the context of transcriptome coordinates since intronic coordinates must be ignored when scanning upstream and downstream [12].

Disregarding some of the mutational events can lead to the identification of different (incorrect) neoantigen nest, which further spawns incorrect neoepitope candidates. For instance, the omission of insertions and deletions (indels) not only alters the underlying DNA sequence but also disregards critical factors such as variant phasing (which determines whether variants occur on the same haplotype), proximal variants, frameshifts introduced by nearby indels, and the influence of custom gene annotation files. Each of these factors can significantly impact neoepitope prediction, as illustrated in Figures 1b–e.

In this study, we systematically evaluate the effects of variant phasing, proximal variants, and frameshifts caused by proximal indels. However, the influence of gene annotation selection (which affects start codon positioning and reading frame shifts, as shown in Figure 1e) is not assessed, as only a fraction of the resulting neoepitope candidates would be accurately identified.

To address these challenges, we introduce Integral Neoantigen Analysis with the Entirety of Mutational Events (INAEME), a comprehensive framework that integrates all key mutational events relevant to neoantigen prediction. Table 1 lists the supported features and mutational events in publicly available Neoantigen workflows and INAEME, our proposed solution. The influence of indels, proximal variants, frameshifts of proximal indels, and variant phasing on obtained neoepitope candidates will be measured in the Results section. The possibility to include in the analysis the custom gene annotation file with alternate splicings is available only in INAEME, while CloudNeo searches for the start codon (AUG) in the protein sequence, and other solutions use transcripts obtained by variant prediction effect tools. The possibility to run the entire analysis from raw reads to scored neoantigens and description of the workflow enable an easier and more flexible user experience, but, in essence, does not change the final neoantigen sequence so they won’t be included in the measurement tasks. RNA expression and variant calling can significantly contribute to the filtering of the final list of neoepitope candidates but their influence will not be measured in this manuscript due to a lack of matching RNA data. The variability in the genome introduced by structural variants and alternate splicing will not be considered here since their accurate and reliable detection is not yet available in NGS data.

**Table 1:** List of neoantigen detection features or mutational events present in four publicly available algorithms (pVACSeq, CloudNeo, PGV, PVC) and our proposed solution (INAEME).

| Feature or mutational event | INAEME | pVACSeq | CloudNeo | PGV | PVC |
| --- | --- | --- | --- | --- | --- |
| Indels | + | + | + | + |  |
| Proximal (neighboring) variants | + |  |  | + | + |
| Frameshift created by proximal indels | + |  |  |  |  |
| Variant phasing | + |  |  | + | + |
| Custom gene/transcript annotations, excluding introns, backward and forward strands | + |  |  |  |  |
| Integral analysis (from FASTQ to scored neoantigens) | + |  |  | + | + |
| RNA expression | + | + |  | + |  |
| RNA variant calling | + |  |  |  |  |
| Described in a workflow description language | + |  | + | + |  |

## Results

To demonstrate the benefits of considering the entirety of mutational events (alternations of the reference genome and thus, the proteins caused by gene variants produced from the individual’s genome), we processed 300 samples with the INAEME neoantigen discovery workflow that originate from The Cancer Genome Atlas (TCGA) [16], containing tumor and normal DNA reads (Supplementary Table 1). Additionally, to better understand the sensitivity of INAEME to each of the included mutational parameters, seven different configurations of INAEME were run - each configuration included a different combination of mutational events. In total, 2400 analyses (tasks) were executed on the Cancer Genomics Cloud platform [17]. The execution time of the neoantigen discovery workflow on an AWS computational cloud instance (c5.4xlarge - 16 processors and 32 GB of memory) varied between 2 and 12 hours per sample, depending on the cancer type and mutational burden (Figure 2 in Supplementary). The samples used in the analysis are selected from three different tumor types to match the benchmarking dataset for the PVC toolset [12]: 300 total cases comprising 100 cases each of melanoma, hepatocellular carcinoma, and lung squamous cell carcinoma. On these selected samples, we measured the differences in the obtained neoantigen nests and neoepitopes when some of the mutational events were not considered. Mutational events investigated here include indel variants, the influence of proximal indels, phasing, and window size around variants to look for proximal variants. We also measured the influence of ignoring one or more mutational events in current publicly available neoantigen discovery algorithms. We made INAEME flexible to allow “switching off” the selected mutational events to measure the exact consequence of neglecting one or more of them, as in the publicly available algorithms. The consequence of neglecting mutational events is the appearance of falsely detected (FD) and falsely excluded (FE) neoantigen nests and neoepitopes. In the rest of the paper, we will use FD and FE terms to distinguish candidates that were wrongly discovered or incorrectly omitted due to neglecting some of the mutational events. Candidates that appear in both scenarios, with and without the exclusion of mutational event(s), we will call correctly detected (CD). Figure 2a shows neoantigen nests comparisons across three cancer types, the counts of correctly detected and the distributions of FD and FE appeared when one or more of these mutational events are switched off, respectively:

1. All indels are ignored,
2. The proximal area size of only 89 nucleotides, as it is used in PVC,
3. All proximal variants are ignored (the proximal area around the variant equals zero),
4. Excluding variant phasing (phasing off), and
5. Influence of indels around the currently processed somatic variant.

**Figure 2:**
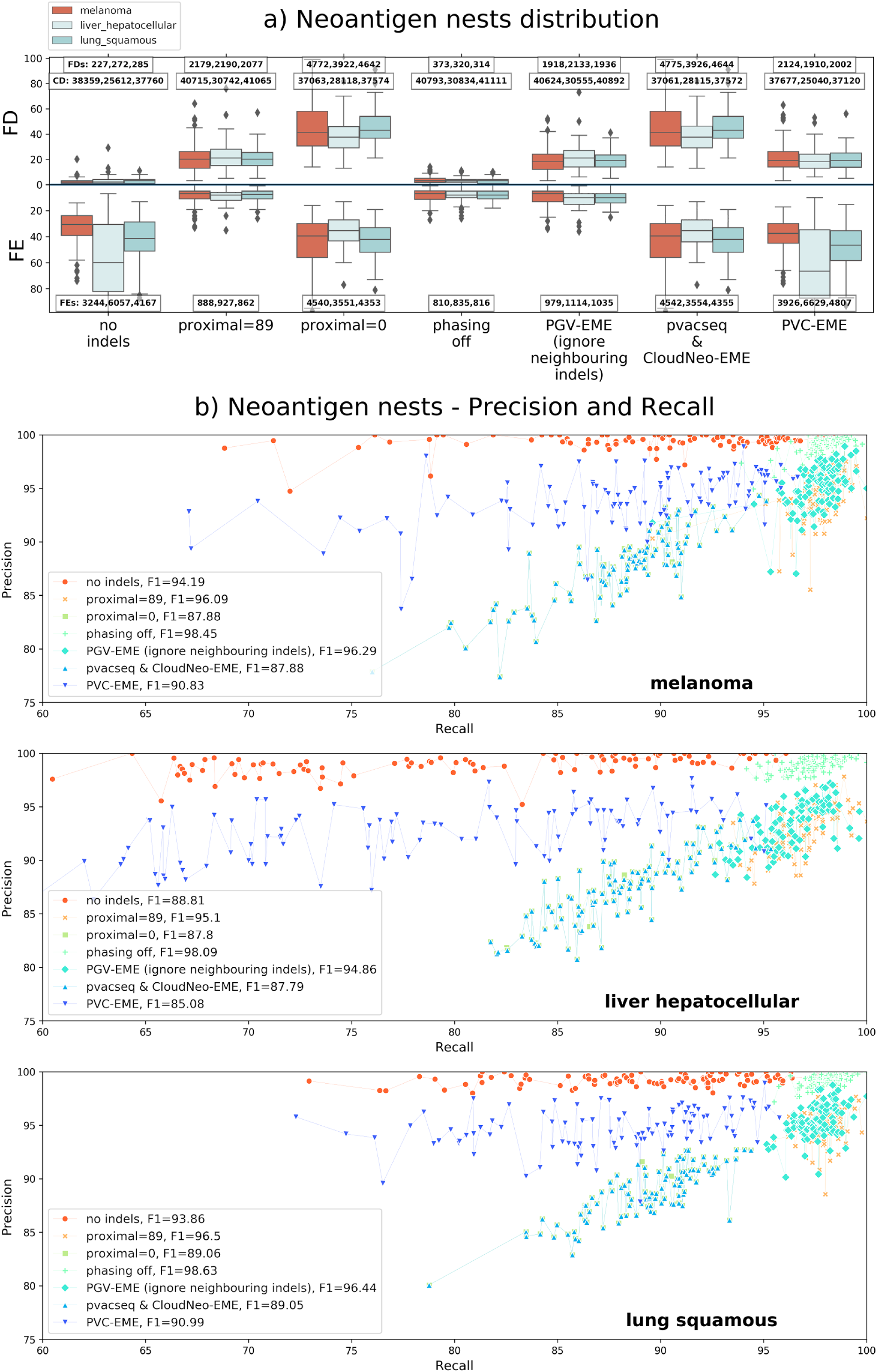
(a) The distribution of falsely discovered (FD), falsely excluded (FE), and correctly detected (CD) neoantigen nests across 100 samples of melanoma, hepatocellular carcinoma, and lung squamous cancer is presented with a mirrored y-axis for better comparison. To identify the influence of including the mutational events on the final list of identified neoantigens, some of the mutational events are sequentially switched off (ignored) for comparison, as presented on the x-axis. The influence of the following mutational events is tested: ignoring all indels, using the proximal area size of only 89 nucleotides as used in PVC, ignoring all proximal variants (proximal=0), and excluding variant phasing (phasing off). Additionally, results of different algorithms are compared on the same graph: PGV - PGV-EME (excluding the influence of indels around currently processed somatic variant), pvacseq & CloudNeo - EME (excluding phasing and proximal area equals 0), and PVC-EME (no indels and proximal area of 89). The y-axis shows the total number of FDs (upper diagram) and the total number of FEs (lower diagram) per sample. Total numbers of FDs, FEs, and CDs for all samples are given in the boxes per cancer type for each mutational event switched off and for each algorithm with its mutational events switched off. To obtain an easier comparison between the distributions of FD and FE neoepitopes, we mirrored the y-axis in the diagram. (b) Precision and recall scores on neoantigen nests across 100 samples each of melanoma, hepatocellular carcinoma, and lung squamous cancer when the same mutational events are switched off. Mean f-scores (F1) across all samples when a mutational event is switched off are given in the figure’s legend.

The effect of phasing, as depicted in Figure 1a, appears lower compared to proximal = 0 because phasing primarily resolves the haplotype context of variants, whereas setting proximal = 0 excludes neighboring variants, including indels that introduce frameshifts, which can lead to entirely different amino acid sequences. As illustrated in Figure 1b, phasing information may, in some cases, produce the same peptide sequence as when phasing is absent, whereas disregarding proximal variants results in a greater number of sequence alterations. This is particularly relevant for cases where indels induce frameshifts, causing substantial deviations in predicted neoepitopes. Consequently, the overall impact of proximal = 0 is more pronounced than the effect of phasing alone.

Figure 2a also shows the comparison of INAEME when all mutational events are considered with INAEME with excluded mutational events (EME) as in publicly available algorithms: PGV (excludes ignore neighboring indels - PGV-EME), pvacseq, and CloudNeo (both exclude phasing and all neighboring variants - pvacseq&CloudNeo-EME), and PVC (excludes indels and has a proximal area of 89 nucleotides - PVC-EME). Figure 2b shows precision, recall, and f-scores for the same listed mutational events exclusion and algorithms, also compared to the algorithm where the entirety of mutational events is taken into consideration. FD, FE, and CD distributions and precision, recall, and f-score for neoepitopes are very similar to the neoantigen nests’ distribution, which makes sense since they are derived from them. They are given in Figure 1 of the supplementary material for completeness of the reported data. An interesting finding here is To assess the impact of falsely identified and excluded neoantigen candidates, we evaluated their major histocompatibility complex (MHC) binding affinity, which is a key determinant of whether a neoepitope is presented on the surface of antigen-presenting cells. Figure 3 illustrates the contribution of different mutational events to the loss of potentially strong neoantigens.

**Figure 3:**
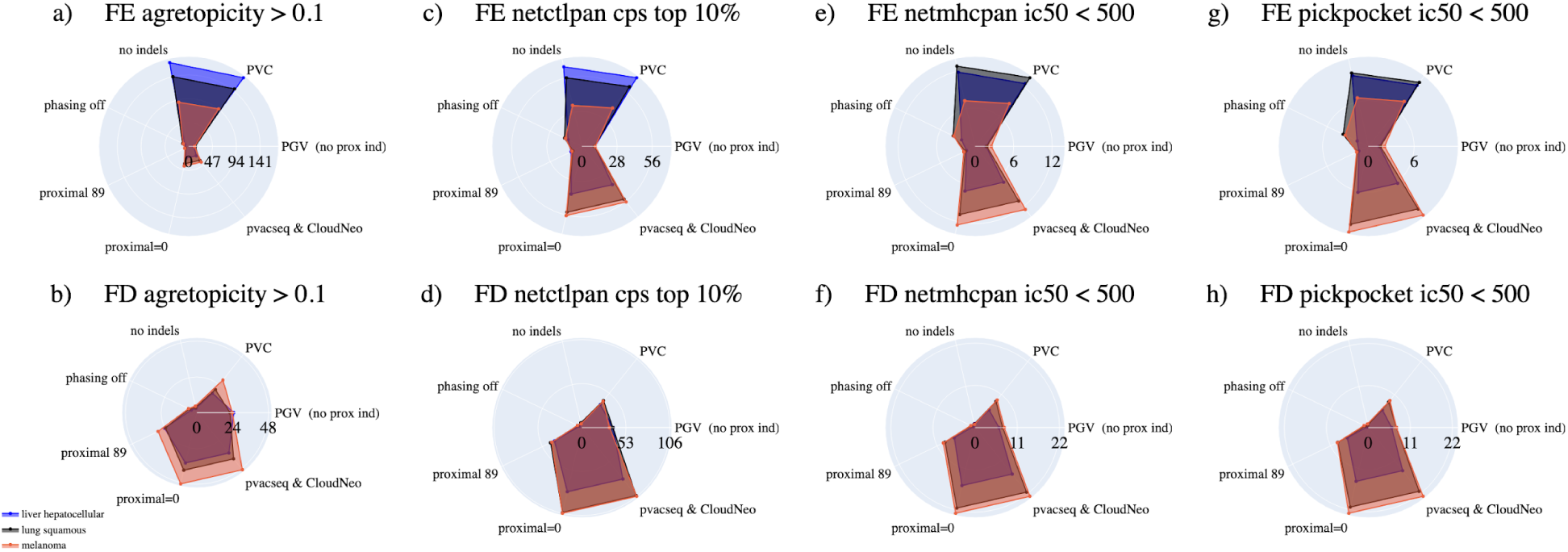
Mean per sample counts of strong neoantigen candidates across 300 TCGA samples of 3 cancer types for falsely excluded and falsely discovered neoantigen candidates, with the exclusion of the single mutational events and exclusion of the mutational events from publicly available algorithms (radial axis). Mean count per sample of FE (a) and FD (b) neoantigen candidates with agretopicity larger than 0.1 calculated by the NetCTLPan tool. A total of 327 neoantigen candidates on average per sample with agretopicity > 0.1 were called without the exclusion of any mutational event. Mean per sample count of FE (c) and FD (d) candidates within the top 10% NetCTLPan’s combined prediction scores. A total of 565 neoantigen candidates on average per sample that were within the top 10% scored by NetCTLPan, are called without the exclusion of any mutational event. Mean per sample count of FE (e) and FD (f) candidates with ic50 < 500 calculated by NetMHCpan. A total of 114 neoantigen candidates on average per sample were with ic50<500 scored by NetMHCpan and called without exclusion of any mutational event. Mean per sample count of FE (g) and FD (h) candidates with ic50 < 500 calculated by PickPocket. A total of 112 neoantigen candidates on average per sample were with ic50<500 scored by PickPocket and called without exclusion of any mutational event.

We applied three bioinformatics tools implemented in INAEME to quantify this effect. First, we identified candidates with an agretopicity score (the difference in MHC binding affinity between the mutated and corresponding reference peptide) greater than 0.1, as calculated by NetCTLPan [28], a threshold indicative of significant binding alterations. The mean number of such candidates per sample from falsely discovered (FD) and falsely excluded (FE) groups, stratified by cancer type and mutational event, is shown in Figure 3a–b. To further assess the impact of incorrect variant handling, we examined FD and FE neoantigen candidates ranked in the top 10% of all predicted candidates by NetCTLPan (Figure 3c–d). Additionally, we analyzed FD and FE candidates with an IC50 binding affinity below 500 nM - considered a strong binding threshold [32] - as predicted by NetMHCpan [29] and PickPocket [30] (Figure 3e–h). This analysis provides insights into how different mutational events affect the identification of high-confidence neoantigens. Across all four selection criteria for strong neoantigen candidates, falsely excluded (FE) candidates predominantly arise from the omission of indels, whereas falsely discovered (FD) candidates are primarily generated by disregarding proximal variants.

The largest stumbling block in the neoantigen discovery area is the lack of truthset samples with validated neoantigen candidates. Without a validated truth set, researchers are constrained to make relative comparisons between configurations as we’ve done in Figs 2-3. To move towards a fair comparison between neoantigen discovery algorithms, here we show the impact of selected mutational events and highlight the necessity for their inclusion in the neoantigen discovery algorithms. The first initiative to create a truth set and propose best practices in neoantigen discovery has been started through the Tumor Neoantigen Selection Alliance (TESLA) challenges organized by the Parker Institute for Cancer Immunotherapy. INAEME was evaluated across three rounds of the TESLA challenges (Rounds 2, X, and 3). When applied to eight tumor samples from [19], comprising 38 neoantigen candidates validated by microscopy and flow cytometry, INAEME successfully identified and highly ranked 34 of the 38 confirmed neoantigen candidates (Figure 4).

**Figure 4:**
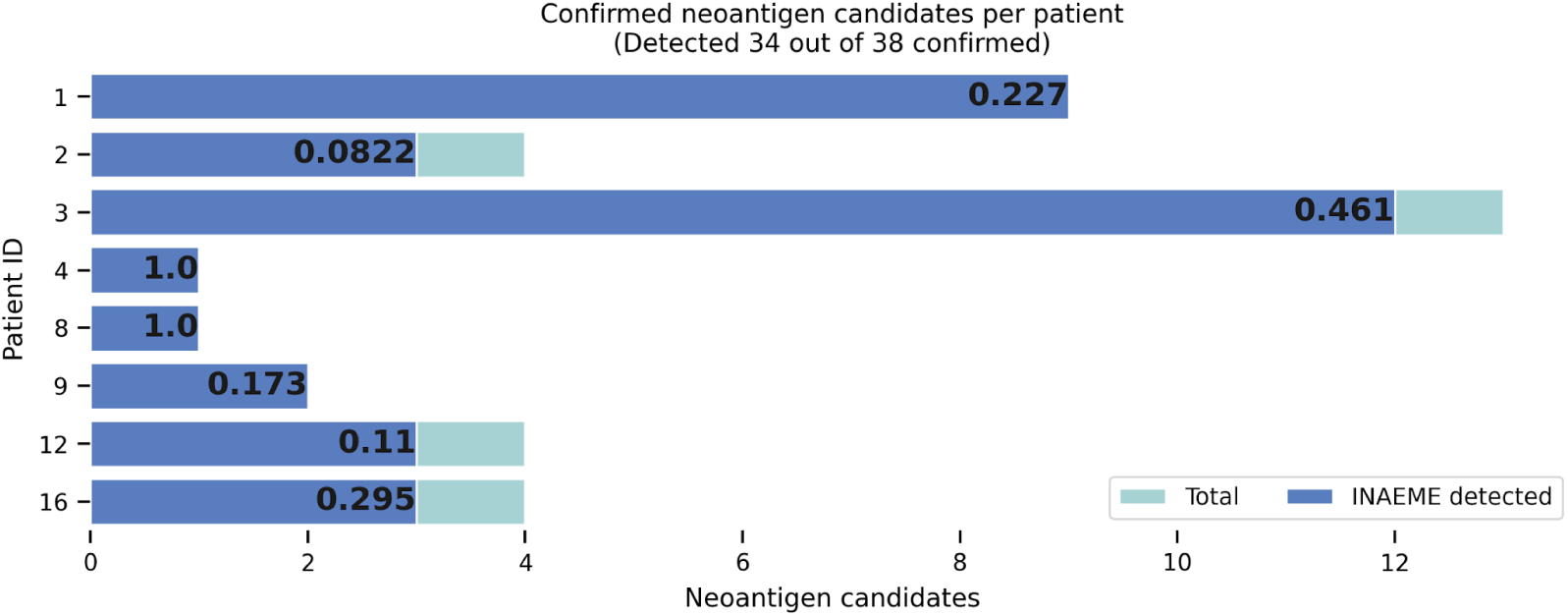
Flow cytometry and microscopy confirmed neoantigen candidates from [19] detected and scored by INAEME with ranked AUPRC metric across 8 patients. INAEME final neoantigen prediction score was created by combining NetMHCPan IC50 score with a weight of 0.7 and RNA expression score with a weight of 0.3, which was validated throughout rounds of TESLA challenges. Neoantigen candidates were also filtered with the following criteria: NetCTLPan combined prediction score > 0.2 and tumor VAF > 0.1.

To assess INAEME’s ability to prioritize immunogenic neoantigens, the ranked area under the precision-recall curve (AUPRC) was used, a metric that evaluates the ranking of true neoantigens relative to non-immunogenic candidates [19]. AUPRC provides a fair comparison by accounting for the relative position of each confirmed neoantigen among all predicted candidates. A perfect AUPRC score of 1 is achieved when all validated neoantigens are ranked highest, whereas the presence of high-ranking false positives lowers the score. For patients 4 and 8, each had a single confirmed neoantigen, which INAEME ranked as the top candidate, yielding an AUPRC of 1 for both cases.

Still, we were unable to analyze the four missed candidates in detail since the publicly available TESLA data does not contain the individual validation details they provided during the challenges. As a result, we could not directly evaluate how other tools performed on this same subset of validated neoantigens. Future studies may be able to address this limitation.

## Discussion

The main idea behind developing and testing INAEME is to highlight and prove the significance of including comprehensive mutational events. Neglecting some strong-MHC-binding neoantigen candidates leads to lowering the chance of constructing an adequate cancer vaccine. Cancer vaccines with a smaller number of neoantigens might not be as effective due to the mutational frequency of the tumor and the possible elimination of immunogenic tumor clones [33]. On the other hand, including some of the false neoantigen candidates, besides not targeting cancerous cells for destruction, increases the risk of provoking hazardous autoimmune reactions [34]. Our tests across 300 TCGA samples demonstrated a significant number of falsely called and missed neoantigen nests and candidates when we intentionally excluded some of the mutational events. As a summary, we compared INAEME (implemented with the entirety of mutational events) with the INAEME version with excluded mutational events from publicly available neoantigen discovery algorithms such as PGV, pvacseq, CloudNeo, and PVC. Our neoantigen discovery analysis showed 384 neoantigen nests (putative sources of neoantigen candidates) on average per sample. Across 300 TCGA samples, we obtained 20 (5.2%) incorrect and 10 missed (2.6%) neoantigen nests for PGV [11] on average per sample. For PVACseq [9] and CloudNeo [10], 44 (11.45%) were incorrect, and 42 (10.93%) were missed, and for PVC [12], 20 (5.2%) were incorrect, and 51 (13.28%) were missed (detailed distributions per cancer type are shown in Figure 2). We also tested MHC binding affinities for missed and falsely called neoantigen candidates and showed the existence of strong neoantigen candidates among them (Figure 3).

When comparing distributions of missed neoantigen nests between these 3 cancer types, the noticeable difference when ignoring proximal variants (proximal=0) is around 1000 more FE neoantigen nests across all melanoma and lung squamous samples compared to the liver hepatocellular samples; this could be explained by the larger mutational burden present in these two cancer types [34].

Minimizing both FDs and FEs neoantigen nests is very important: FEs to avoid elimination of potentially correct neoantigen candidates and FDs to remove incorrect, potentially hazardous neoantigen candidates. From our results, the mutational event with the highest influence on FEs is ignoring indels, and it eliminates, on average, 32.44 (8.45%) correct neoantigen nests on melanoma (f-score: 94.19), 60.57 (23.65%) on liver hepatocellular with f-score 88.81, and 41.67 (11.03%) on lung squamous with f-score 93.86 samples. The significance of indels in generating strong neoantigen candidates is proven in many studies, such as [33], where results show that indels generate three times as many predicted neoantigen candidates as non-synonymous single-nucleotide variants and nine times as many strong neoantigen candidates. A smaller number of indels than expected on the melanoma samples is due to a smaller proportion of indels in this cancer type. This result matches with findings in [33], where melanoma has the second smallest proportion of indels out of a total of 19 analyzed cancer types.

The second mutational event that highly influences the appearance of FEs is excluding proximal variants, which create, on average, 45.40 (12.25%) FEs in melanoma, 35.51 (12.63%) in the liver, and 43.53 (11.58%) in lung cancer samples. Ignoring proximal variants also creates the largest number of FDs across all 3 cancer types: 47.72 (12.87%) on average per sample for melanoma, 39.22 (13.95%) on the liver, and 46.42 (12.35%) on lung cancer, with average f-scores per sample, respectively, 87.88, 87.8, and 89.05. Both indels and proximal variants contribute to or create open reading frames that might be an ideal source of tumor-derived neoantigens and thus induce multiple neoantigen-reactive T cells [33]. From all observed mutational events phasing has the smallest, but still significant influence on both FEs (8.10 on melanoma, 8.35 on the liver, and 8.16 on lung on average per sample) and FDs (3.73 on melanoma, 3.20 on the liver, and 3.14 on lung cancer type on average per sample) and with the average per sample f-scores respectively 96.9, 95.1 and 96.5. It is worth noticing that the number of FDs and FEs, while ignoring the proximal variants, has the largest fluctuation between samples, from 10 to 125, and neglecting indels has a large fluctuation of FEs (10 to 125) with f-scores spanning between samples from 60 to 97 (Figure 2b).

Another strong motivation for developing INAEME is to build an integral analysis for neoantigen discovery, starting from raw sequenced reads to neoantigen candidates with all important features that contribute to prioritizing and selecting the best ones. For that reason, each produced neoantigen candidate, besides peptide sequence and HLA type, contains features such as the information about the DNA somatic variant from which it originates including tumor and normal variant allele frequencies, MHC binding prediction scores (NetCTLPan, NetMHCPan, Pickpocket), agretopicity, RNA expression of the transcript from which is extracted, RNA variant and its allelic count. Additionally, neoantigen candidates present in wildtype protein databases are optionally filtered out. All filtering parameters and settings are flexible and exposed in the CWL description of the workflow, making them easy to configure before starting the execution of the workflow.

Accurate, reproducible, and rapid discovery of neoantigen candidates represents a foundation for personalized cancer vaccine development. Paired with a cloud environment and integral analysis entirely described in CWL, the INAEME workflow has the potential to fulfill the demands for rapid turnaround of accurate predictions within a clinically compliant environment for a large number of patients. Although developed in the cloud environment, sometimes due to data usage restrictions and privacy policy, it is not able to process samples in the cloud. Still, it is possible to run INAEME on the computing cluster or any machine using the Rabix executor (https://github.com/rabix/bunny).

Future work on this topic should focus on improvements of the workflow. In the first place, adding support for MHC Class II binding prediction tools, to be able to capture and process the full spectrum of the neoantigen candidates. Additional confirmation of the quality of the produced neoantigen candidates currently existent as the limitation of this study should be done through the new comparison to other existing solutions..

## Methods

INAEME is described in Common Workflow Language [18], where all of the total 130 bioinformatics tools are executed inside corresponding Docker containers, making it fully reproducible and portable to any computational platform. It is also highly configurable, having a number of exposed parameters that enable simple comparison analysis and measuring of the influence of certain settings. The execution flow of INAEME is similar to the general neoantigen discovery flow [20][21] with some additional processing steps (Figure 5).

**Figure 5:**
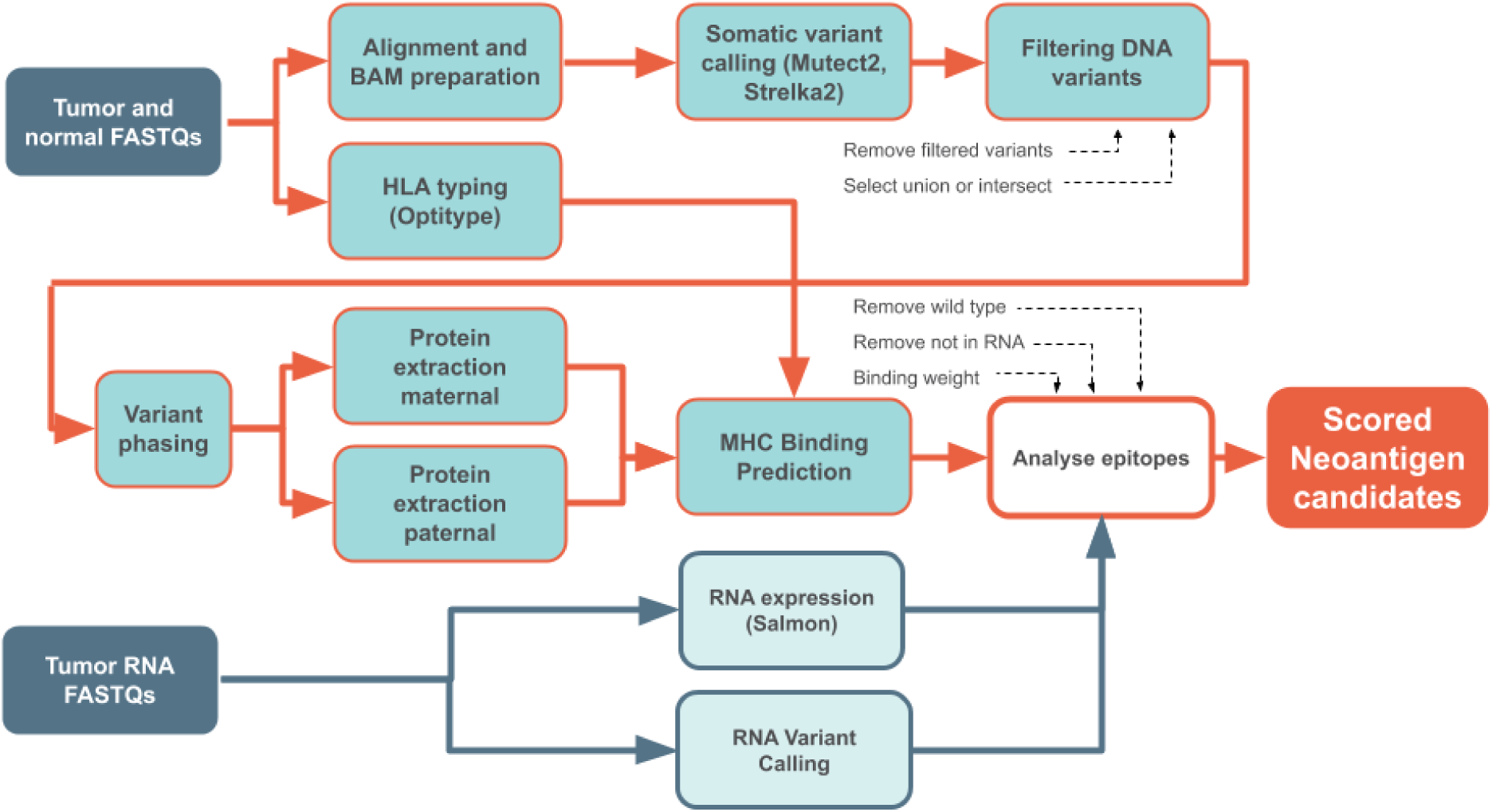
Detailed processing diagram of INAEME.

The first step performs read alignment [23], leveraging GATK best practices. Next, somatic variant calling using Mutect2 [24] and Strelka2 [25] is performed, and variants are merged into one file. Merging is configurable and includes selecting the intersect (default) or the union of variants from these two callers. Once the somatic variants are merged, an optional and recommended step is variant phasing, which identifies alleles on maternal and paternal chromosomes (Figure 1b) [26]. The protein extraction step runs in two instances, called maternal and paternal, separating variants with the same locus coming from the different haplotypes. It is in charge of reconstructing the neoantigen amino acid sequence by taking into account all of the mentioned mutational events.

In parallel with the Alignment step, the HLA typing step is performed by running the OptiType [27] tool to calculate Human Leukocyte Antigen (HLA) types from tumor DNA FASTQs, which will be later used for the MHC Binding Prediction step.

In parallel with DNA-seq analysis, RNA transcript expression quantification is performed to enable the selection of influential DNA variants that will be translated into proteins of interest. RNA reads are processed with Salmon and/or RSEM tools, which enable high-precision quantification of transcripts. The result of RNA expression analysis is a list of gene isoforms with corresponding RNA expression scores. RNA reads are used to call RNA variants within the RNA Variant Calling step.

Once DNA, RNA, and HLA processing steps are finished, the data obtained are used to determine potential neoantigens and their ranking based on HLA compatibility. To do so, phased somatic DNA variants, optionally filtered with RNA variants (if RNA data is available to boost the precision of neoantigen candidates), are passed to the Analyse Epitopes tool. This step consists of several custom-developed Python tools that apply all variants to the reference genome and translate the extracted nucleotide sequences around the somatic variants into proteins (neoantigen nests). The area around the altered amino acid that defines the neoantigen nest is configurable and set by default to 10 amino acids on both sides, giving, for a single nucleotide variant, a total size of 21 amino acids. The area of 10 amino acids around the altered amino acid is chosen because the maximal peptide size of MHC Class I presented antigens on the surface of the cell is 10 (the groove will accommodate peptides of approximately 8-10 amino acids in length [37]), and for the same reason, the maximal size for neoantigen candidates supported in our binding prediction tools is 11. With that, the altered amino acid will be included in all neoantigen nests extracted from that neoantigen nest. Protein Extraction Tools take into account the full range of above mentioned mutational events, currently not existent within the published neoantigen workflows, and also: changes from both germline and somatic variants (SNPs and indels), variants within the STOP codon (nonstop), and mutations that create STOP codons (nonsense). The extracted protein sequence and identified HLA type are used by the epitope prediction tools NetCTLpan [28], NetMHCpan [29], and PickPocket [30], trained on peptides from the Immune Epitope Database (IEDB) [31] to compute confidence-ranked neoantigen candidates.

Finally, a list of the confidence-ranked neoantigen candidates with HLA types, variant information, RNA expression, and RNA variant calling scores is merged and prioritized in the Analyse epitopes tool. The final score on which neoantigen candidates are prioritized is configurable and, by default, calculated using a combination of normalized epitope binding score from the MHC binding prediction tool (selectable from the list of tools - NetCTLPan, NetMHCPan, Pickpocket) and normalized RNA expression. The weight parameter applicable to these two scores is also configurable. Our experiments conducted on samples from the Tesla challenges [19] with available sets of flow-cytometry validated neoantigen candidates showed that the weighting normalized MHC binding score with 0.7 and normalized RNA expression with 0.3 gives the best final score (more trust is given to the MHC-binding score). Despite that, these weights are not universal and might depend on many factors such as the quality of the MHC binding prediction tools, the quality and depth of RNA sequencing data, the specific cancer type being analyzed, and the experimental validation methods used. The transcript to which the DNA somatic variant falls can be highly expressed, which would give neoantigen candidates originating from it a very high score, but this score is incorrect if the reference allele of the transcript is expressed and not the alternative (mutated) one. To mitigate this scenario, we added a configurable possibility enabled by RNA variant calling to filter false-positive neoantigen candidates originating from DNA variants that are not called in RNA. Measuring the influence of RNA variant calling on obtaining accurate neoantigen candidates is not considered in this paper and represents one of the targets for future work.

Agretopicity of neoantigen candidates is also one of the features calculated in the Analyse epitopes tool together with tumor and normal VAFs (variant allele frequencies). If a wild-type protein database is provided to Analyse epitopes, it is possible to additionally filter neoantigen candidates that are identical to some wild-type protein. For example, some variants could be falsely called somatic and, with available wild-type databases, they could be filtered out. Additionally, neoantigen candidates containing three or more repeated amino acids in a row are filtered out by default. MHC molecules present peptides to T cells based on their ability to fit within the binding groove. Peptides with low sequence complexity, such as those containing long stretches of the same amino acid, often lack the structural diversity required for stable and high-affinity binding. The anchor residues that facilitate binding typically rely on a specific pattern of hydrophobic, polar, or charged residues, which may be disrupted by excessive repetition of a single amino acid.

The Analyse epitopes tool produces the final report table with prioritized neoantigen candidates (peptide-HLA pair) and all above-mentioned features and scores for it (originating somatic DNA variant location, filter and info fields, scores from all MHC-binding tools, RNA expression, allelic depth and depth of originating RNA variant if called, tumor and normal VAFs, agretopicity, etc). In that way, the user might select different criteria to prioritize neoantigen candidates.

## Data availability

The data used in this paper are available in the Cancer Genome Atlas (TCGA) database with sample IDs given in Supplementary Table 1.

## Code availability

The INAEME workflow described in CWL is available for use in academic research projects without restrictions on the Cancer Genomics Cloud platform, and it is available for download as a BioCompute object (JSON-based standards for describing computational workflows in genomics): https://biocomputeobject.org/builder?https://biocomputeobject.org/BCO_000462/DRAFT. CWL description of the workflow with source code of all scripts and command lines for all bioinformatic tools are given in the supplementary material of this manuscript (*INAEME.json*).

## User’s guidelines

INAEME is an end-to-end workflow for discovery and prioritization of Class I neoantigens from tumor and normal samples. It integrates DNA-seq and RNA-seq analysis, HLA typing, variant phasing, peptide reconstruction and MHC class I binding prediction to produce a ranked peptide–HLA candidate list with rich annotations.

The most convenient way to run the workflow is through the Cancer Genomics Cloud platform, with available free access and funds. After registering, the user should:

1. Create the project
2. Create a new workflow
3. In the code field, paste the *INAEME.json, save, and run the workflow*
4. In the draft task page, provide necessary inputs, including raw FASTQ files for tumor and normal samples, reference genome, and gene annotations
5. After all inputs are set, run the task Here are the example of input files:

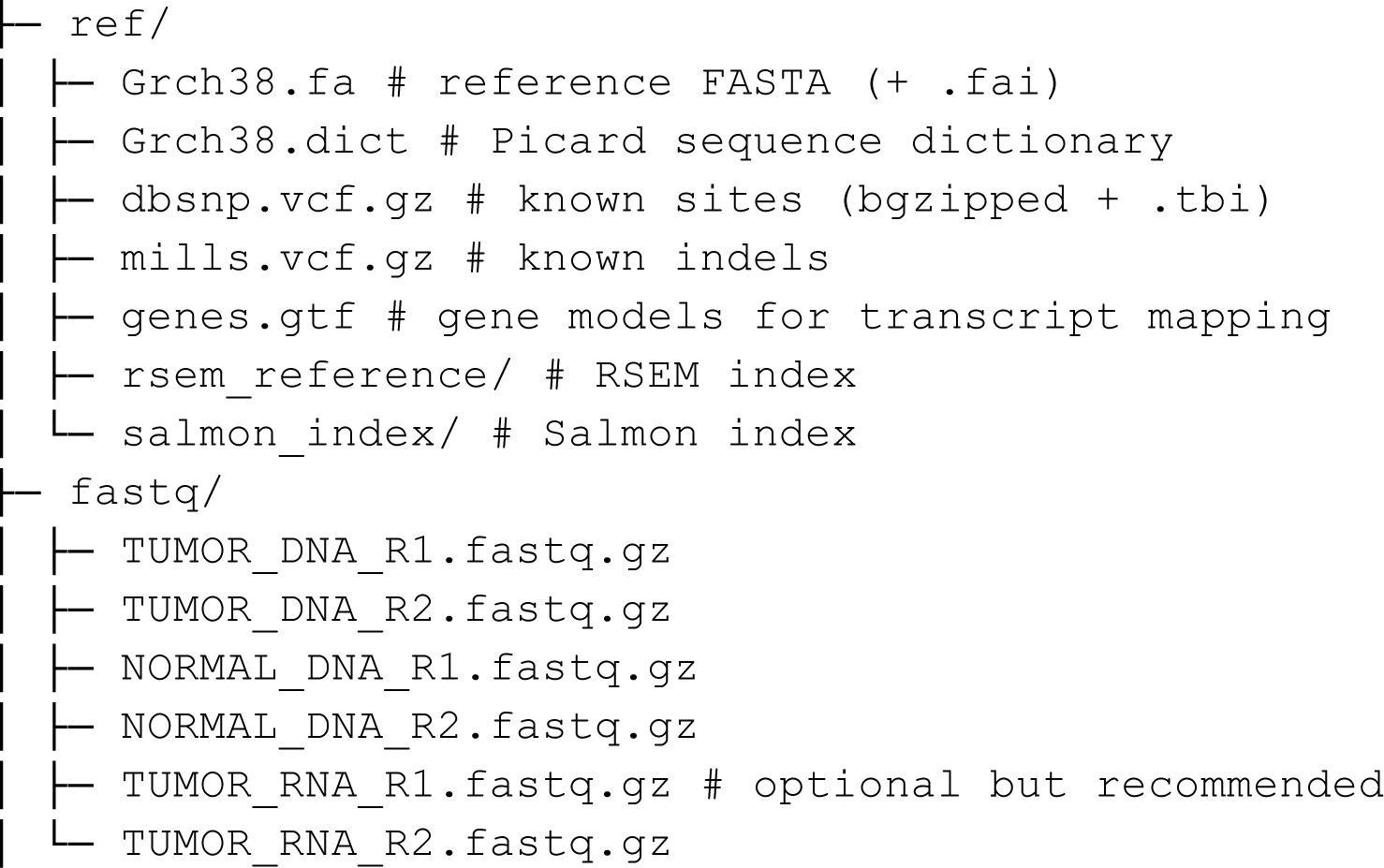

## Acknowledgments

The authors thank Mladen Lazarevic, Brandi Davis-Dusenbery, Aleksandra Kovacevic, Nevena Veljkovic, Julie Gil, and Jeffrey Brabec for reviewing the paper and providing useful comments.

The results published here are in whole or in part based upon data generated by the TCGA Research Network: https://www.cancer.gov/tcga. dbGaP access for processing TCGA samples was enabled by the National Institutes (request #88805-1 for access phs000178/GRU, INC5023870).

## Author contributions

V. K. was involved in all aspects of this study, including constructing and developing

The initial idea, methodology, analyzing and interpreting data, and writing the manuscript. O. M. contributed to the development of the initial idea, methodology for variant translation, protein extraction, and visualizations. N.I.R. and M. K. were developing the RNA analysis part of the workflow and helped with the review of the paper. A.M.L., N.K., and J.D. were involved in project management, overseeing the progress, discussions, and review. All authors reviewed the manuscript.

## Supplementary

**Figure 1:**
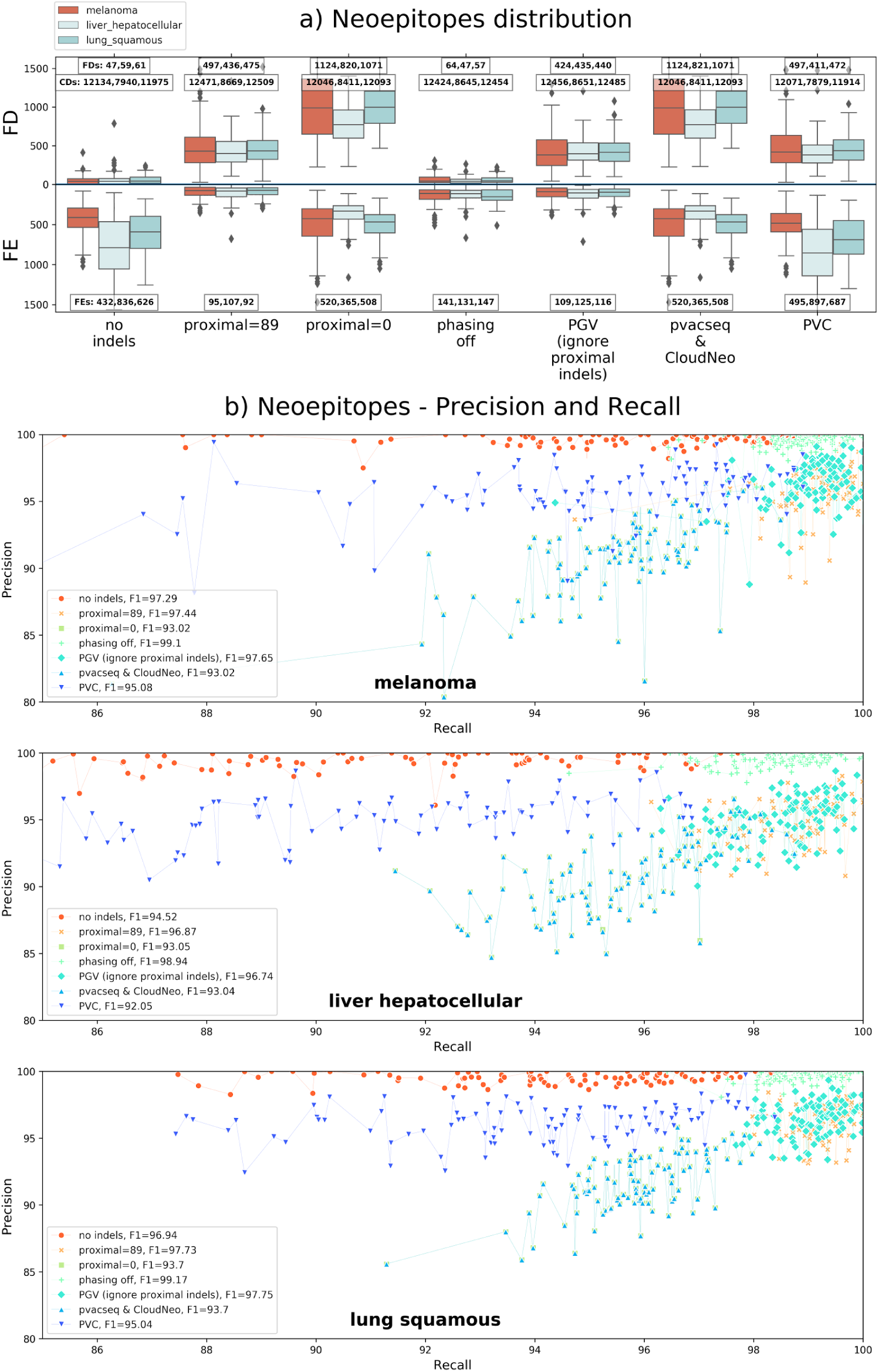
(a) Distribution of FDs, CDs, and FEs on neoepitopes across 100 samples of melanoma, hepatocellular carcinoma, and lung squamous cancer, respectably, when some of the mutational events are switched off: all indels are ignored, the proximal area size of only 89 nucleotides as used in PVC, all proximal variants ignored (proximal=0), excluding variant phasing, the influence of indels around currently processed somatic variant (as excluded in PGV), pvacseq & CloudNeo (excluding phasing and proximal area equals 0) and PVC (no indels and proximal area of 89). Total numbers of FDs, FEs, and CDs are given in the boxes for each mutational event switched off and for each algorithm with its mutational events switched off. (b) Precision and recall scores on neoepitopes across 100 samples each of melanoma, hepatocellular carcinoma, and lung squamous cancer when the same mutational events are switched off. Mean f-scores (F1) across all samples when a mutational event is switched off are given in the figure’s legend. Neoepitopes (sizes 8, 9, 10, and 11 amino acids) are spawned from all neoantigen nests not contained in the wild-type protein database.

**Figure 2:**
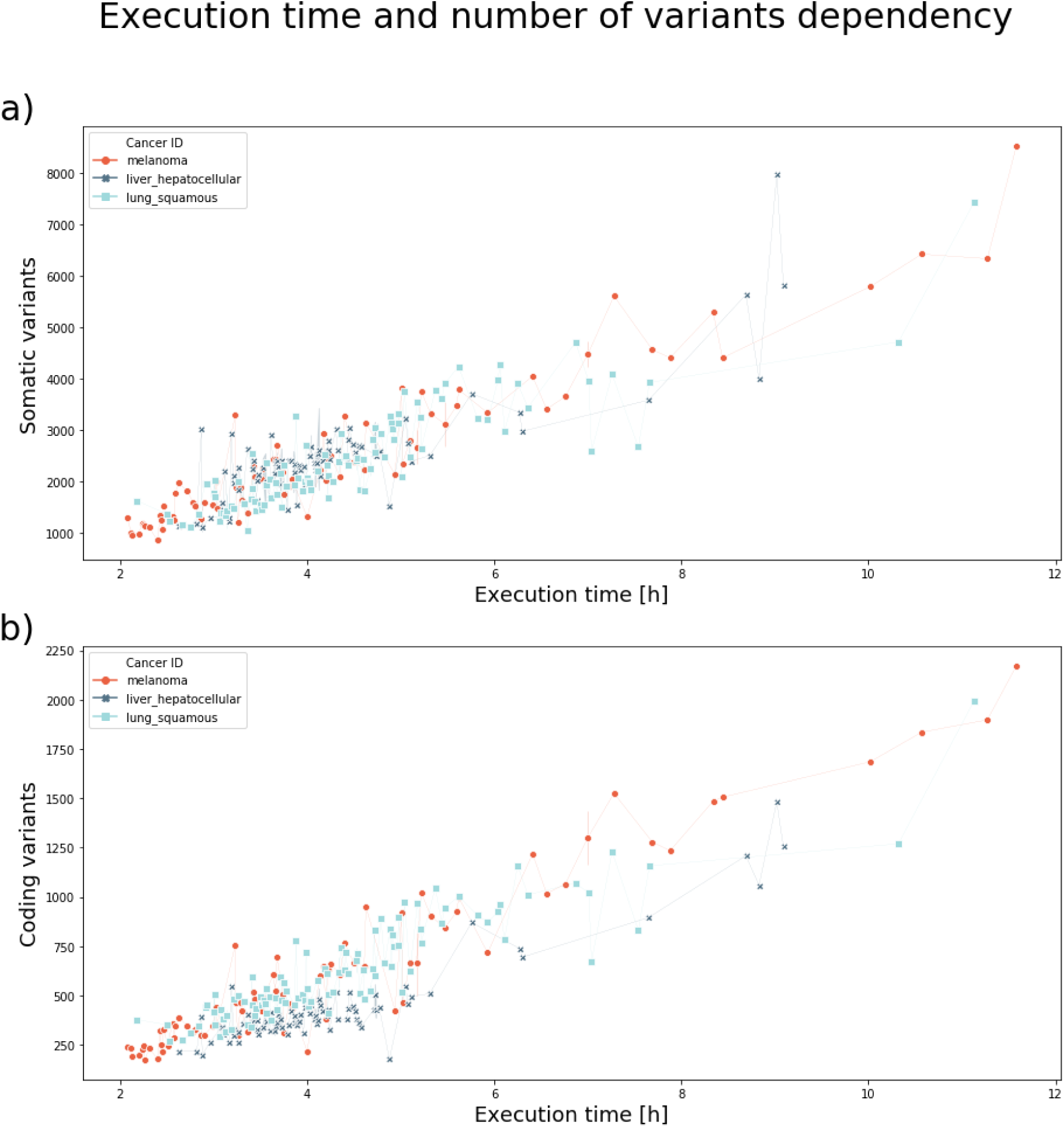
Distribution of INAEME execution times across 100 samples of melanoma, hepatocellular carcinoma, and lung squamous cancer (total of 300 samples). (a) Dependency of execution time and number of somatic variants detected in the sample. (b) Dependency of execution time and number of somatic variants that fall into gene coding regions.

**Table 1:** Case IDs of TCGA samples used to analyze mutational events.

