## Supplementary figures and images for "INAEME: Integral Neoantigen Analysis with Entirety of Mutational Events"

### sup_fig1_300_epitopes.png

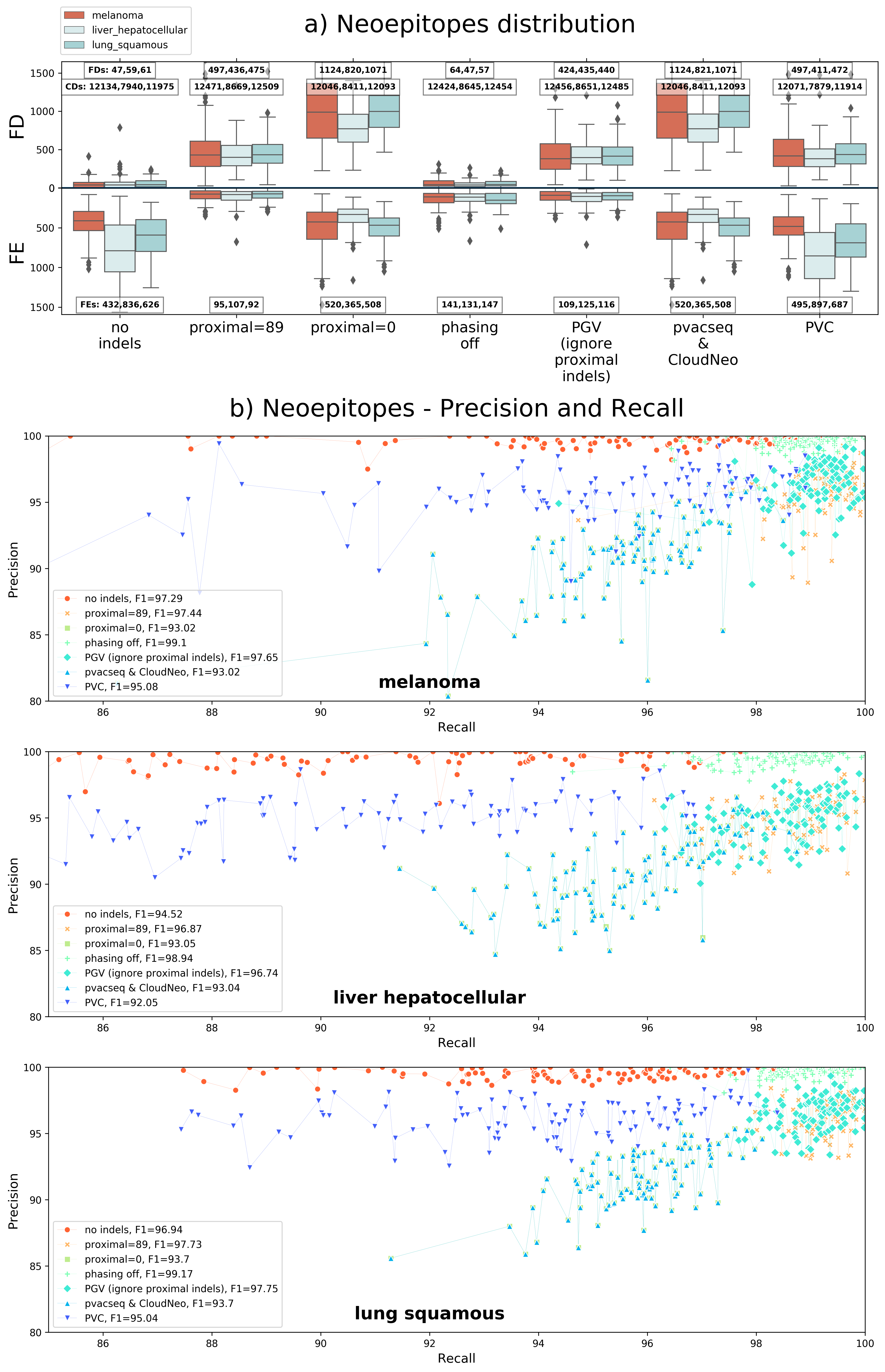

### sup_fig2_execution_time.png

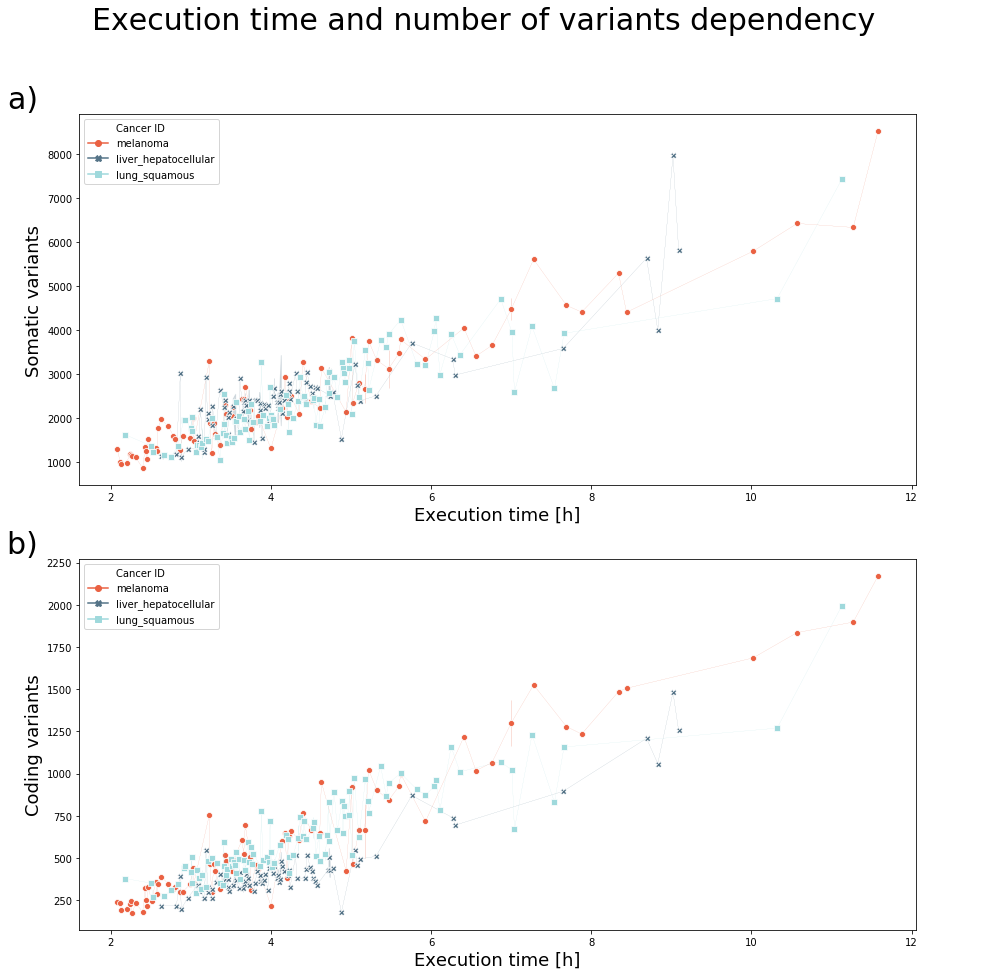
